# Parallel comparison of pre-conditioning and post-conditioning effects in human cancers and keratinocytes upon acute gamma irradiation

**DOI:** 10.1101/450106

**Authors:** Jason Cohen, Nguyen T. K. Vo, Colin B. Seymour, Carmel E. Mothersill

## Abstract

**PURPOSE:** To determine and compare the effects of pre-conditioning and post-conditioning towards gamma radiation responses in human cancer cells and keratinocytes

**MATERIALS AND METHODS:** The clonogenic survival of glioblastoma cells (T98G), keratinocytes (HaCaT), and colorectal carcinoma cells (HCT116 p53^+/+^ and p53^-/-^) was assessed following gamma ray exposure from a Cs-137 source. The priming dose preceded the challenge dose in pre-conditioning whereas the priming dose followed the challenge dose in post-conditioning. The priming dose was either 5 mGy or 0.1 Gy. The challenge dose was 0.5 – 5 Gy.

**RESULTS:** In both pre- and post-conditioning where the priming dose was 0.1 Gy and the challenge dose was 4 Gy, RAR developed in T98G but not in HaCaT cells. In HCT116 p53^+/+^, pre-conditioning had either no effect or a radiosensitizing effect and whereas post-conditioning induced either radiosensitizing or radioadaptive effect. The different observed outcomes were dependent on dose, the time interval between the priming and challenge dose, and the time before the first irradiation. Post-conditioning effects could occur with a priming dose as low as 5 mGy in HCT116 p53^+/+^ cells. When HCT116 cells had no p53 protein expression, the radiosensitizing or radioadaptive response by the conditioning effect was abolished.

**CONCLUSIONS:** The results suggest that radiation conditioning responses are complex and depend on at least the following factors: the magnitude of priming/challenge dose, the time interval between priming and challenge dose, p53 status, cell seeding time prior to the first radiation treatment. This work is the first parallel comparison demonstrating the potential outcomes of pre- and post-conditioning in different human cell types using environmentally and medically relevant radiation doses.

## INTRODUCTION

In biology and medicine, pre-conditioning refers to exposing organisms or cells to a large damage-inducing dose of a stressor preceded by a smaller non-toxic dose of the same stressor whereas in post-conditioning a large dose of the stressor is followed by a smaller dose (Calabrese et al., 2007). The conditioning effect is generally built upon the hormesis concept (Calabrese and Mattson, 2017). In principle, conditioning enables the activation or enhancement of protective programs and therefore helps reduce the damaging effect caused by the stressor (Calabrese et al., 2007; Calabrese and Mattson, 2017). The conditioning effect is perhaps best studied in vascular biology. Ischemic conditioning contributes to cardioprotection by reducing oxidative stress and thus minimizing or preventing myocardial infarction (Aimo et al., 2015).

The concept of pre-conditioning also applies to radiation biology where radiation is the stressor. Radiation pre-conditioning describes the irradiation of subjects to a large damage-causing dose after a smaller dose (known as priming dose), subsequently leading to a more radio-resistant phenotype (Wolff 1998). Olivieri et al. (1984) were the first group to report the protective effect of radiation preconditioning in human lymphocytes exposed to low-dose radioactive thymidine. Radiation preconditioning, now widely known as “radioadaptive response” (RAR), is observed with different types of radiation and in many biological systems (Azzam et al., 1994; de Toledo et al., 2006; Maguire et al., 2007; Ryan et al., 2008; 2009; Day et al., 2006; Smith and Raaphorst, 2003; Audette-Stuart and Yankovich, 2011; Tang et al., 2016; Vo et al., 2017a; Smith et al., 2011). In cancer onset, radiation preconditioning can extend the latent period (Lemon et al., 2017b). The protective effect can occur with the priming dose of as low as 0.001 mGy (Day et al., 2006). The mechanism governing radiation pre-conditioning or RAR appears to evolutionarily conserved (Calabrese and Mattson, 2017). As life evolved with radiation for millions of years, this concept largely contributes to the big picture of adaptation, longevity, and evolution of biological entities living in environments with low radioactivity as well as their interactions with other bystander participants within the ecosystem. In medicine, cancer patients are subjected to diagnostic imaging procedures (low dose exposure) prior to an actual therapeutic radiation regimen (high dose exposure). Thus the pre-conditioning-induced radio-resistance may reduce the effectiveness of the intended radiation therapy program.

Radiation post-conditioning, on the contrary, is the procedure of irradiating subjects with a large dose before a very small priming dose. To date, the literature on radiation post-conditioning is relatively small with significantly fewer studies (Day et al., 2007; Lemon et al., 2017a; Lin and Wu, 2015). In general, radiation post-conditioning exerts a similar RAR-like effect as seen in pre-conditioning (Day et al. 2007; Lemon et al., 2017a). The protective effects of radiation post-conditioning include enhanced protection against chromosomal damage (Day et al., 2007) and increased lifespan (Lemon et al., 2017a). The lowest priming dose inducing a RAR in radiation post-conditioning recorded to date was 0.01 mGy (Day et al., 2007). Currently, how comparable the extent of radioprotection is between preconditioning and post-conditioning is not well understood. We previously observed that a smaller dose preceding a large dose had a less damaging effect than the reverse order (Mothersill and Seymour, 1993), suggesting that different mechanisms might underlie low and high dose response and that irrespective of the order of delivery both mechanisms were activated by experience of different doses. The data also suggested immediate and delayed death levels involved different mechanisms. Lin and Wu (2015) later showed that differences in effects were cell type-and species-dependent in a complex multi-dose fractionated radiation setup. Interestingly, in one of their two-dose experiments with a human colon cancer cell line HT-29, post-conditioning had a more protective effect than preconditioning.

Increasing evidence has suggested that there are a number of factors that are associated with or directly involved in RAR, especially in the context of low-dose radiobiology. Nitric oxide radicals and other reactive oxygen species are key players mediating the RAR (Matsumoto et al. 2007; Miller et al., 2016; Murley et al., 2017; 2018). More interestingly, factors in the irradiated cell conditioned medium can alleviate some of the damages caused by high dose in bystander cells (Maguire et al., 2007; Matsumoto et al., 2000; 2001). Cells that can develop RAR also have traits of hyper-radiosensitivity/increased radioresistance (HRS/IRS) (Ryan et al. 2009). Additionally, the tumour suppressor p53 protein is important in regulating RAR. While p53 regulates apoptosis in low-dose radiation hypersensitivity (Enns et al., 2004), its expression is suppressed during RAR (Takahashi, 2001), which would be consistent with the protective survival mechanism although sparing potentially damaged cells may have a role in post-radiotherapy tumour recurrence. However, in p53-null or -mutant cells, priming cells with a lower dose before a larger challenge dose makes cells more radiosensitive (the complete opposite to the traditional RAR) (Miller et al., 2016; Murley et al., 2017; 2018). A lot of this knowledge has been gained from the radiation pre-conditioning perspective. Yet nothing is known regarding the other conditioning view, i.e. radiation post-conditioning. Most recent research by our group has evidence suggesting that post-conditioning effects can occur on a quantum biological level (Mothersill et al., 2018), possibly via UV biophoton-mediated mechanisms (Le et al., 2017a).

This current paper shows possible outcomes associated with radiation pre-conditioning and post-conditioning effects in different human cell types.

## MATERIALS AND METHODS

### Cell Cultures

Four human cell lines were used in this study. They were two colorectal cancer lines HCT116 p53^+/+^ and HCT116 p53^-/-^, one brain glioblastoma multiforme cell line T98G, and one spontaneously immortalized untransformed keratinocyte cell line HaCaT.

HCT116 p53^+/+^ had wild-type TP53 gene and (Brattain et al., 1981; Popanda et al., 2000). HCT116 p53^-/-^had both TP53 alleles inactivated via the homologous recombination technology (Bunz et al., 1998) and therefore expressed no p53 protein as confirmed by our group (Mothersill et al., 2011; Le et al., 2017b). HCT116 p53^+/+^ was HRS-positive and HCT116 p53^-/-^ was HRS-negative (Enns et al., 2004). The cell lines were cultured in the Roswell Park Memorial Institute (RPMI)-1640 medium supplemented with 10% fetal bovine serum (FBS), 2 mM L-glutamine, 25 mM HEPES, 100 U/ml penicillin, and 100 µg/ml streptomycin. All tissue culture reagents were obtained from Gibco/Life Technologies (Grand Island, NY), unless otherwise specified. Both cell lines were provided by Dr. Robert Bristow (University Health Network, University of Toronto, Toronto, ON).

T98G had a p53 mutant with a single-nucleotide variant leading to an animo acid switch from Met to Ile at codon 237 (Van Meir et al., 1994). T98G displayed HRS (Short et al 1999). T98G was routinely maintained in the Dulbecco’s modified Eagle medium F12 (DMEM/F12), 10% FBS, 2 mM L-glutamine, 25 mM HEPES, 100 U/ml penicillin, and 100 µg/ml streptomycin. The cell line was provided by Dr. Brian Marples (Department of Radiation Oncology, William Beaumont Hospital, Royal Oak, MI).

HaCaT had a mutated p53 gene with two heterozygous mutations: one with a single-nucleotide variant leading to an amino acid switch from His to Tyr at codon 179 and the second with a double-nucleotide variant leading to an amino acid switch at codon 282 (Lehman et al., 1993). HaCaT displayed no HRS (Ryan et al., 2009). The growth medium for HaCaT was the same growth medium used for HCT116 cells with the addition of 1000 ng/mL cortisol. The HaCaT cell line was kindly provided by Dr. Petro Boukamp (German Cancer Research Center, Heidelberg, Germany).

All cell lines were grown at 37°C in an atmospheric environment equilibrated at 5% CO_2_ and 95% air. Subculturing was performed with trypsin/EDTA as previously described by our group (Vo et al. (2017b) for HCT116 p53^+/+^ and Fernandez-Palomo et al. (2016) for T98G and HaCaT).

### Challenge Irradiation and Clonogenic Assay Technique

The cell survival of the control and the irradiated groups was assessed using the clonogenic assay technique developed by Puck and Marcus (1956). Briefly, adherent cells were detached from the tissue culture surface and dissociated into single cells by trypsin/EDTA along with excessive pipetting. Cell concentrations were determined using a BioRad T20 Automated Cell Counter. In T25 tissue culture flasks (BD Falcon), depending on the plating efficiency of the cell lines and the magnitude of the lethal dose, cultured cells were seeded at densities of 20-60 cells/cm^2^.

Depending on the cell lines, cells were seeded in triplicate flasks for 6h or 24h prior to irradiation challenge. Two non-irradiated controls were used. The first control group, regarded as “0 Gy control” was kept at all times in the incubator. The second control group, regarded as “sham-control” (to control for unknown variations imposed by temperature, transport, and background radiation), was brought to the irradiation facility and was subjected to zero radiation dose. Irradiated flasks were subjected to a first radiation dose, incubated at 37°C for 3-5h, challenged again for a second dose, and finally returned to the 37°C incubator (as illustrated in Figure 1). All flasks were left undisturbed for 9 or 10 days and cell colony survivors (having at least 50 cells) were stained with Carbol Fuschin (Ricca Chemical Company, Alington, TX) and scored. The gamma radiation source was 1 kCi ^137^Cs housed in the Taylor Radiation Source (TRS) suite at McMaster University (Hamilton, ON). Irradiated flasks were 27.1 cm away from the radiation source for the dose rate of 260 mGy/min. Thermoluminescent dosimeters were used to measure radiation doses and to ensure the uniformity of the radiation dose delivery. Three independent experiments were performed.

**Figure 1.**
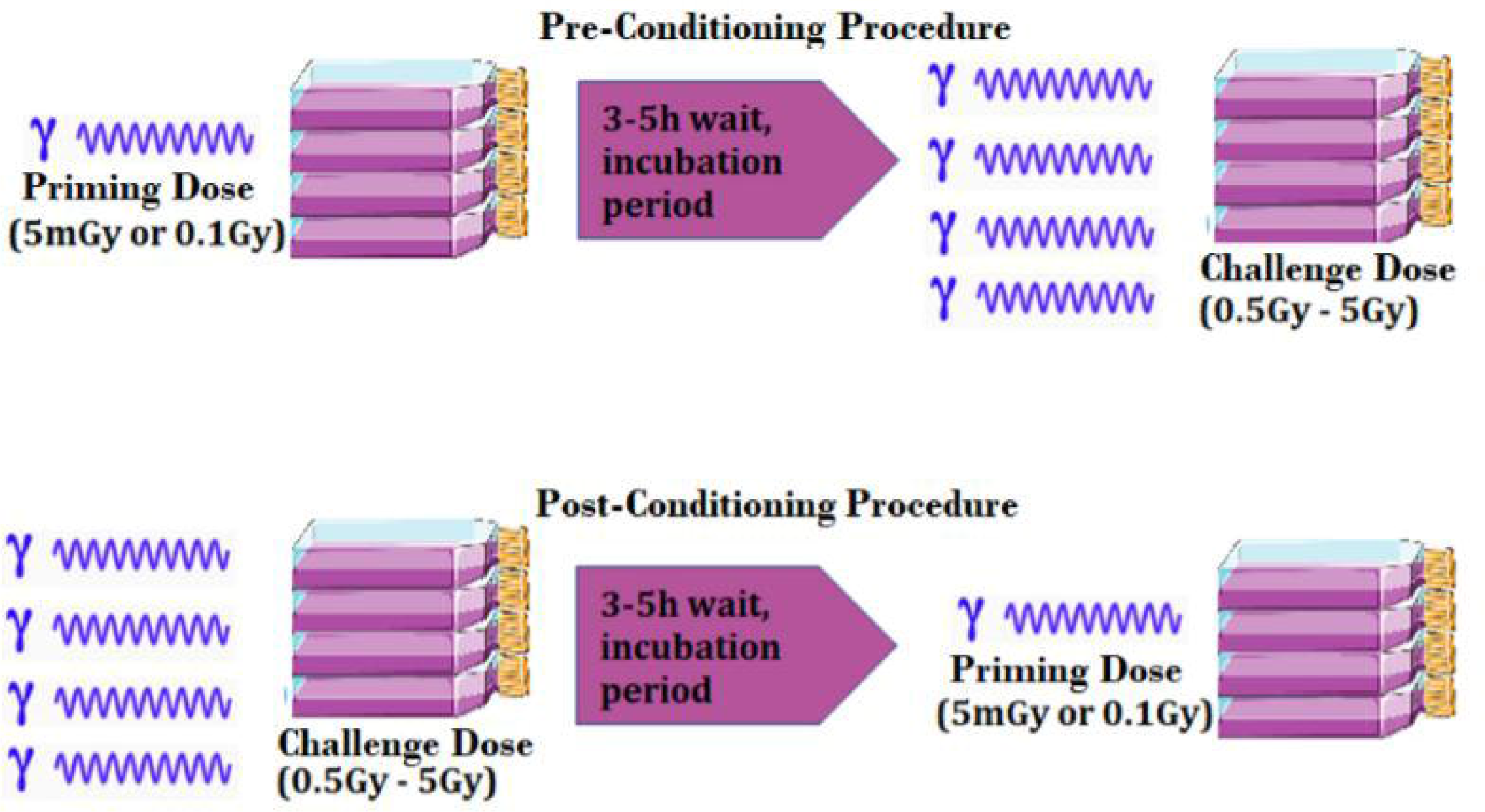
Scheme of the experiments. For both pre-conditioning and post-conditioning treatments, 20-60 cells/cm2 were seeded in triplicate T25 clonogenic flasks. The priming doses were smaller doses and either 5mGy or 0.1 Gy. The challenge doses were larger doses and 0.5-5 Gy. In pre-conditioning, the priming dose preceded the challenge dose. In post-conditioning, the challenge dose preceded the priming dose.

The schemes for the pre-conditioning and post-conditioning response experiments are shown in Figure 1. For the pre-conditioning experiments, the first dose was a small dose (5 mGy or 0.1 Gy) and the second dose was a large dose (0.5-5 Gy). For the postconditioning response experiments, the first dose was a large dose (0.5-5 Gy) and the second dose was a small dose (5 mGy or 0.1 Gy). The small dose of 0.1 Gy was used for all four cell lines while the small dose of 5 mGy was used only for HCT116 p53^+/+^ because (1) the wild-type p53, but not mutant p53, can facilitate RAR (Miller et al., 2016; Murley et al., 2018) and (2) 5 mGy was previously shown to be effective to induce RAR in the HCT116 p53^+/+^ cells (Murley et al., 2017).

### Statistical Analysis

Data are presented mean of clonogenic survival fraction ± SEM (n=9 or n=12). A one-way ANOVA with Tukey’s post test and a confidence interval of 95% or a student’s t-test was used to determine statistical significance via the GraphPad Prism software. p<0.05 were deemed statistically significant.

## RESULTS

### Effects of radiation pre- and post-conditioning in the human glioblastoma cell line T98G

The clonogenic survival was statistically significantly higher in both pre- and post-conditioning than in the no priming dose control (Figure 2). Pre-conditioning and post-conditioning resulted in 7.3 % and 5.3 % increases in clonogenic survival. Therefore, both pre- and post-conditioning induced the RAR in T98G. There was, however, no significant difference in clonogenic survival between pre- and post-conditioning was observed (Figure 2).

**Figure 2.**
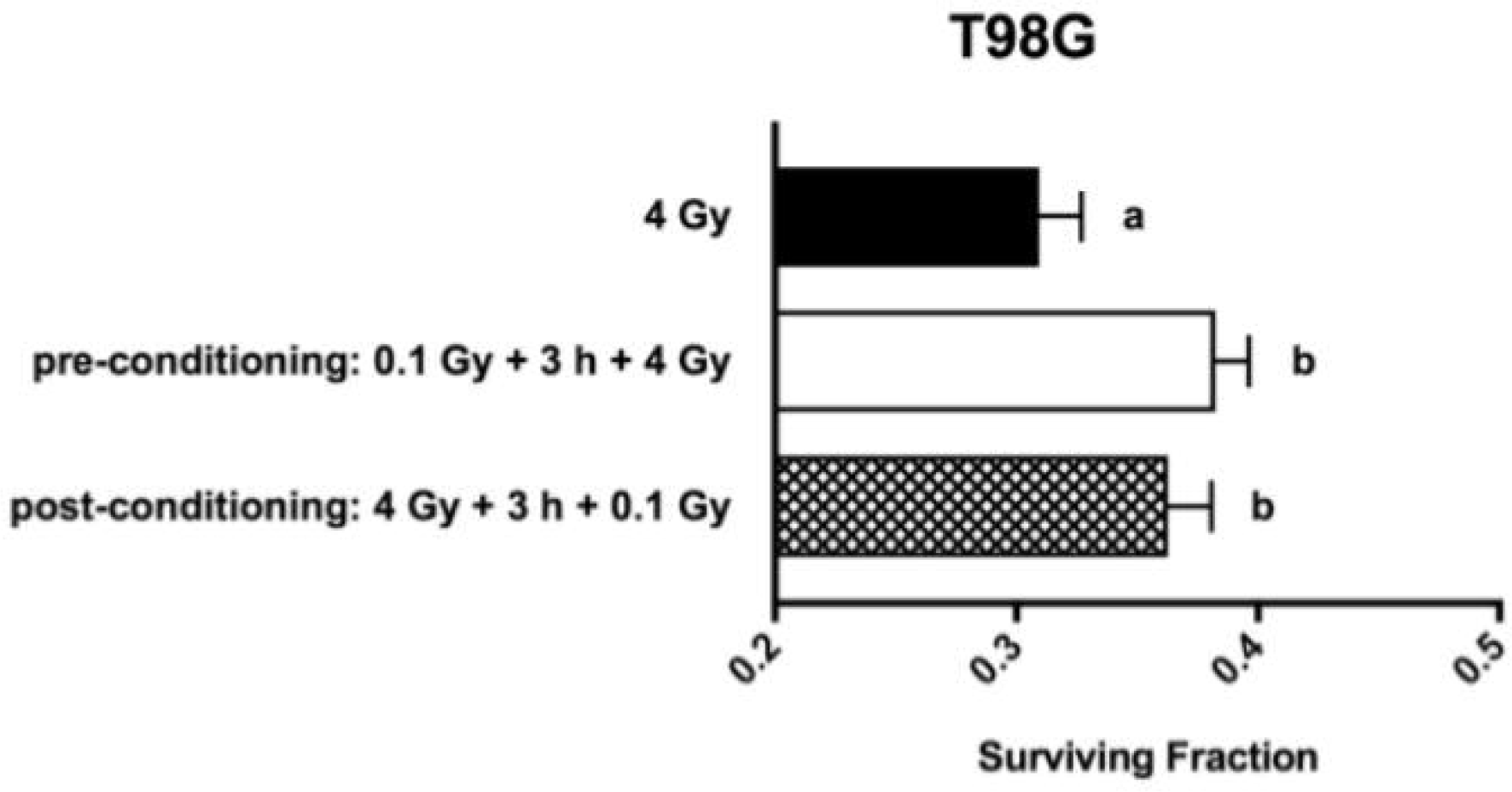
Effects of pre-conditioning and post-conditioning on the clonogenic survival of the human glioblastoma T98G cell line. Irradiation was performed 6 h after seeding cells at 40 cells/cm2 in triplicate T25 flasks. The priming dose was 0.1 Gy and the challenge dose was 4 Gy. The time interval between the priming and challenge dose was 3 h. The data are shown as survival fraction ± SEM (n=9) per treatment group. Based on the one-way ANOVA with Tukey’s post-test, groups with different letters are statistically significant (p<0.05) and groups sharing the same letter are not.

### Effects of radiation pre- and post-conditioning in the human normal keratinocyte cell line HaCaT

The clonogenic survival in pre-conditioning was slightly higher than that of the no priming dose control (Figure 3). In contrast, the clonogenic survival in post-conditioning was slightly lower than that of the no priming dose control (Figure 3). But both cases had no statistical significance. Therefore, no RAR was observed in both pre- and post-conditioning in the HaCaT cells. However, the pre-conditioning clonogenic survival was significantly higher than the post-conditioning clonogenic survival (Figure 3).

**Figure 3.**
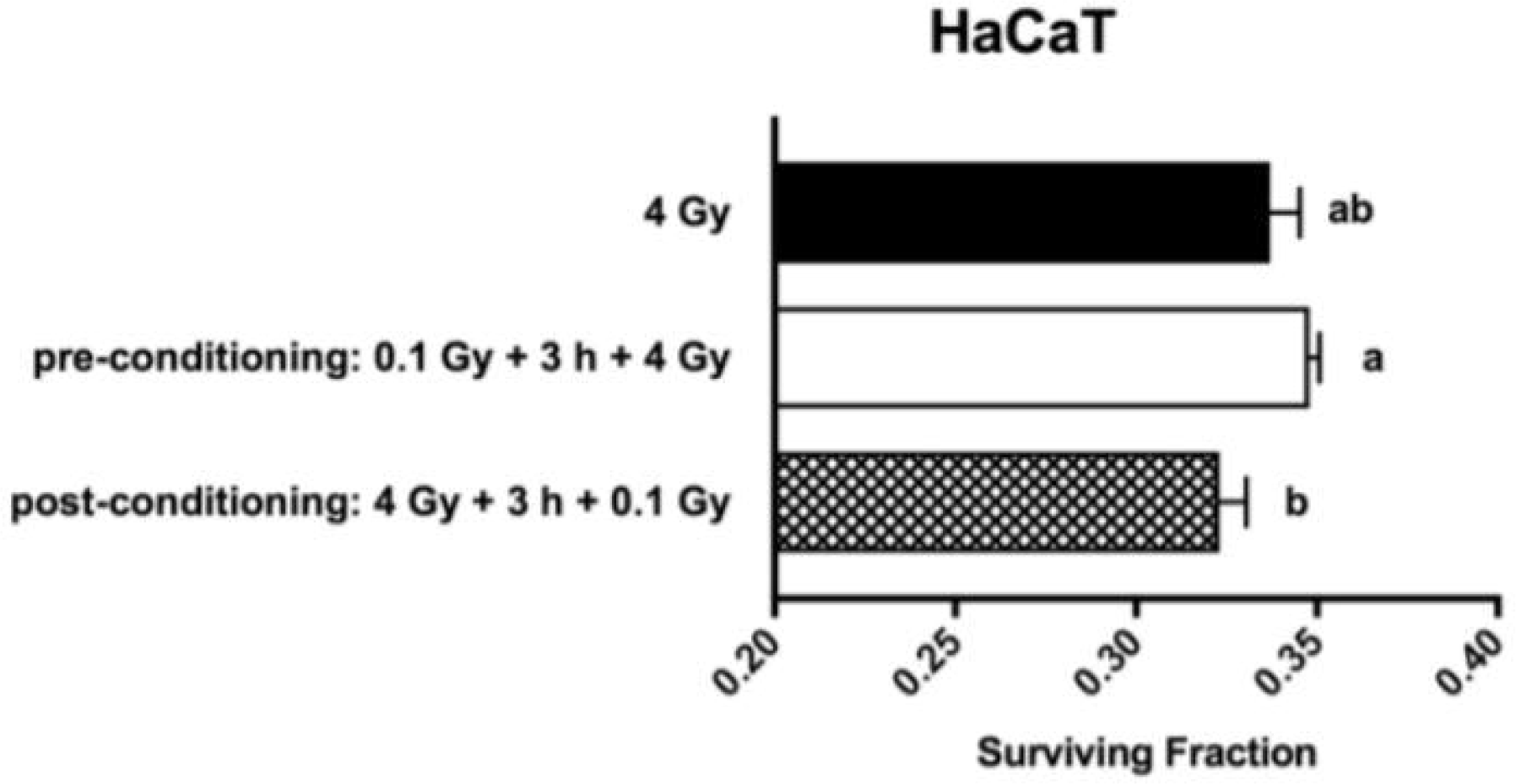
Effects of pre-conditioning and post-conditioning on the clonogenic survival of the normal human keratinocyte HaCaT cell line. Irradiation was performed 6 h after seeding 36 cells/cm2 in triplicate T25 flasks. The priming dose was 0.1 Gy and the challenge dose was 4 Gy. The time interval between the priming and challenge dose was 3 h. The data are shown as survival fraction ± SEM (n=9) per treatment group. Based on the one-way ANOVA with Tukey’s post-test, groups with different letters are statistically significant (p<0.05) and groups sharing the same letter are not.

### Effects of radiation pre- and post-conditioning in the human colon cancer cell line HCT116 p53^+/+^

When the priming dose was 0.1 Gy, the pre-conditioning clonogenic survival was similar to that of the no priming dose control at all three challenge doses (2, 3, and 4) (Figure 4). On the other hand, the post-conditioning clonogenic survival was significantly lower than those in the control and preconditioning at the 2 and 3 Gy challenge doses but not at the 4 Gy challenge dose (Figure 4). Therefore, pre-conditioning did not induce RAR while post-conditioning induced radiosensitization effects in HCT116 p53^+/+^.

**Figure 4.**
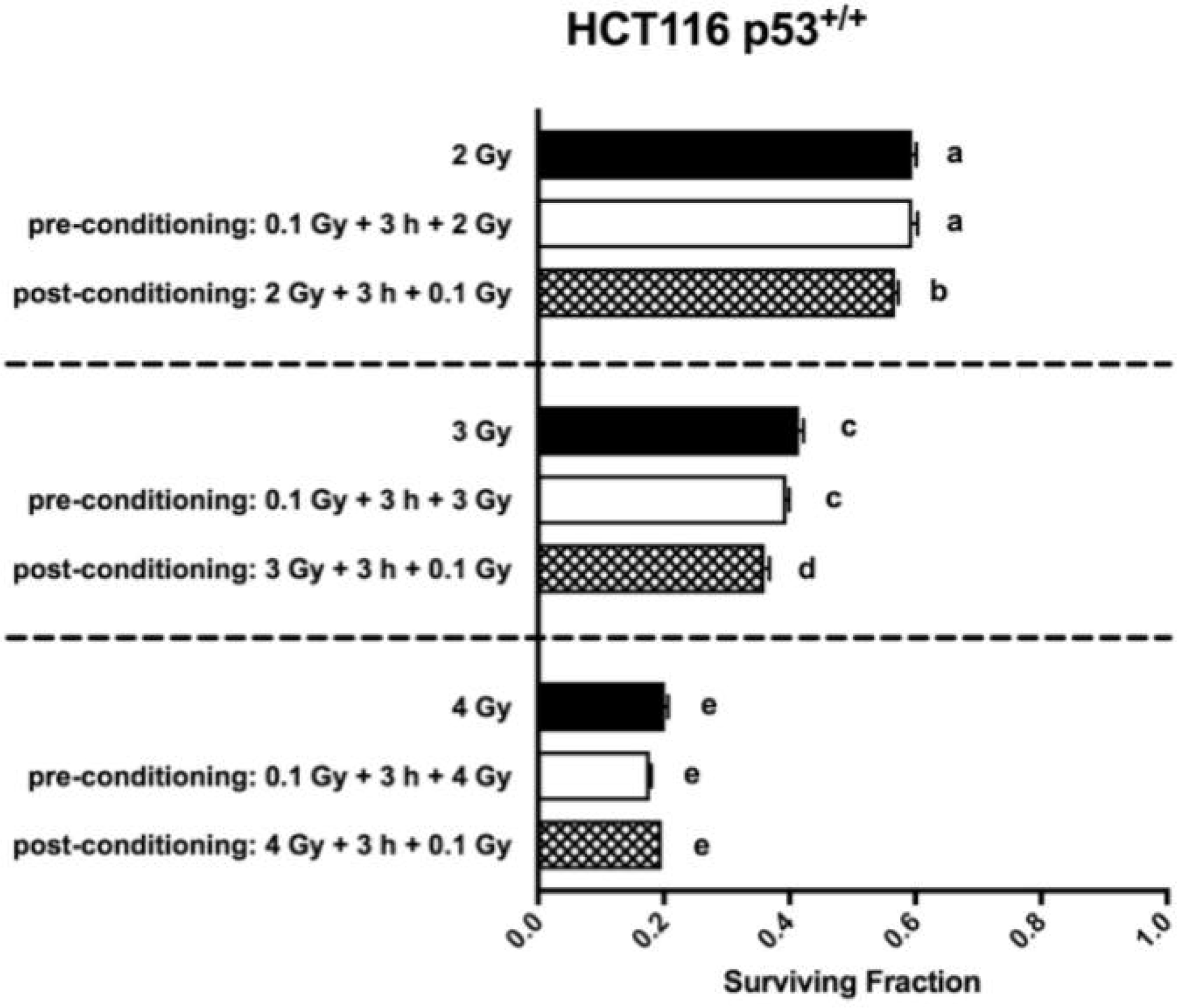
Effects of pre-conditioning and post-conditioning on the clonogenic survival of the human colorectal carcinoma HCT116 p53+/+ cell line. Irradiation was performed 6 h after seeding 40 cells/cm2 in triplicate T25 flasks. The priming dose was 0.1 Gy and the challenge dose was 2, 3 or 4 Gy. The time interval between the priming and challenge dose was 3 h. The data are shown as survival fraction ± SEM (n=9) per treatment group. Based on the one-way ANOVA with Tukey’s post-test, groups with different letters are statistically significant (p<0.05) and groups sharing the same letter are not.

To further see if RAR could be achieved in the HCT116 p53^+/+^ cells at more environmentally relevant doses, the priming dose was chosen to be 5 mGy while the larger challenge dose was 0.5 Gy. In the pre-conditioning, the clonogenic survival was higher (an 5.2 % increase) but not statistically significant than that of the no priming dose control (Figure 5). In the post-conditioning, the clonogenic survival was significantly higher (an 8.4 % increase) than the no priming dose control (Figure 5). However, there was no difference in the clonogenic survival between the pre- and post-conditioning (Figure 5).

**Figure 5.**
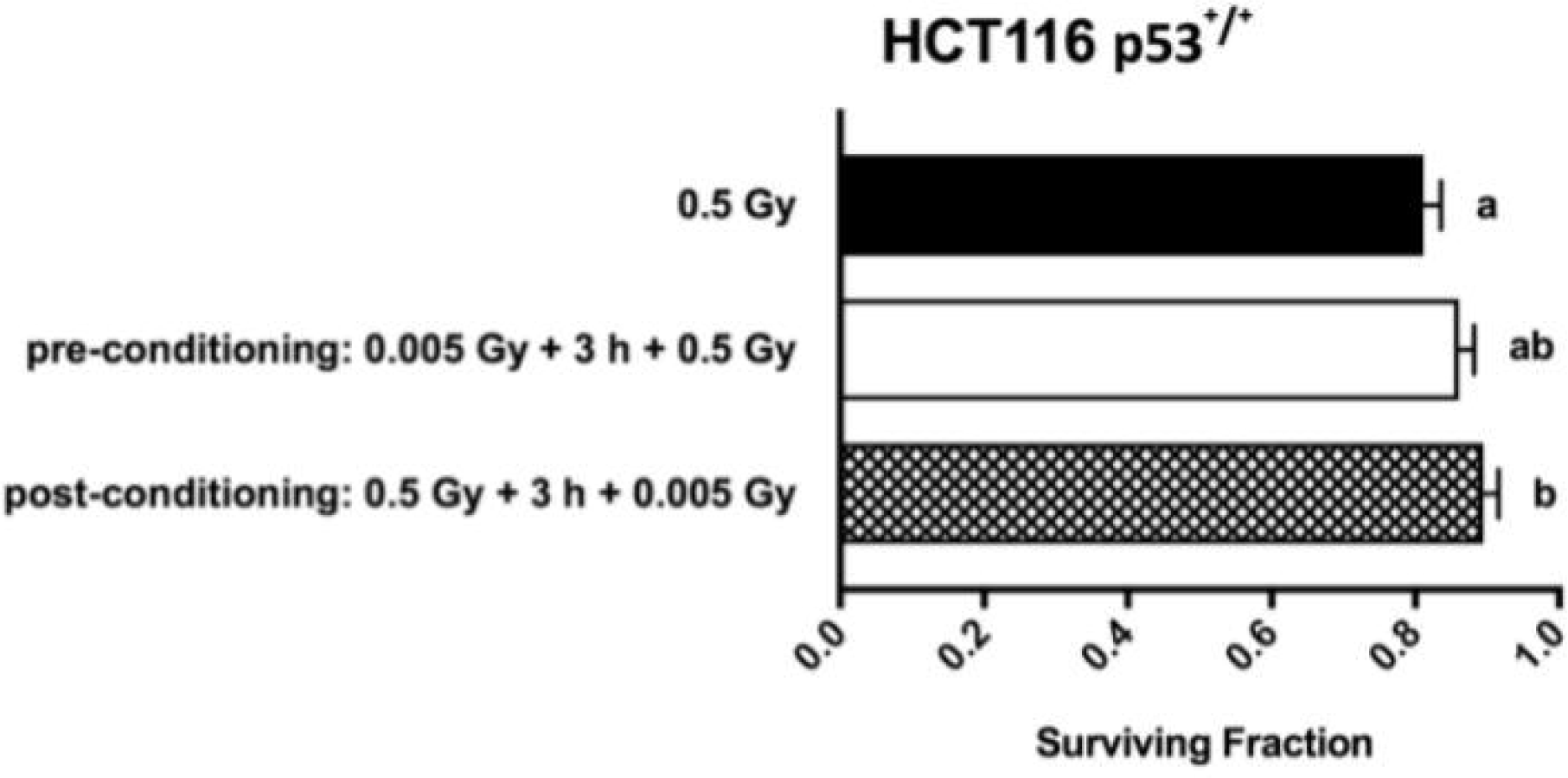
Effects of very low-dose radiation pre-conditioning and post-conditioning on the clonogenic survival of the human colorectal carcinoma HCT116 p53+/+ cell line. Irradiation was performed 6 h after seeding 40 cells/cm2 in triplicate T25 flasks. The priming dose was 5 mGy and the challenge dose was 0.5 Gy. The time interval between the priming and challenge dose was 3 h. The data are shown as survival fraction ± SEM (n=9) per treatment group. Based on the one-way ANOVA with Tukey’s post-test, groups with different letters are statistically significant (p<0.05) and groups sharing the same letter are not.

When HCT116 p53^+/+^ cells were seeded for 24 h and then followed the radiation conditioning and exposure regimen where the time interval between the 0.1 Gy priming dose and the larger challenge dose was extended to 5 h (instead of 3 h in previous experiments), an interesting result was obtained. The no priming dose control for the pre-conditioning (0.1Gy-5h-2Gy) was shown as “0Gy-5h-2Gy” and the no priming dose control for the post-conditioning (2Gy-5h-0.1Gy) was shown as “2Gy-5h-0Gy”. Surprisingly, the clonogenic survival of the no priming dose control for the pre-conditioning was significantly higher than that of the no priming dose control for the pre-conditioning (Figure 6). The pre-conditioning clonogenic survival was significantly lower than its no priming dose counterpart whereas the post-conditioning clonogenic survival was significantly higher than its no priming dose counterpart (Figure 6). Therefore, under these experimental conditions, pre-conditioning sensitized radiation responses whereas post-conditioning induced RAR in HCT116 p53^+/+^ cells. When the TP53 gene coding p53 protein was knocked out in HCT116 p53^+/+^ cells (as in the resulting HCT p53^-/-^ subclone), the clonogenic survival was the same in pre- and post-conditioning as well as in their no priming dose control counterparts (Figure 7A); thus the observed radio-sensitization/adaptive effect was abolished. This was also true when the challenge dose increased to 5 Gy (Figure 7B). This clearly shows that p53 regulated the mechanisms that led to the radiosensitization in pre-conditioning and RAR in post-conditioning in HCT116 p53^+/+^ cells in response to high-dose radiation.

**Figure 6.**
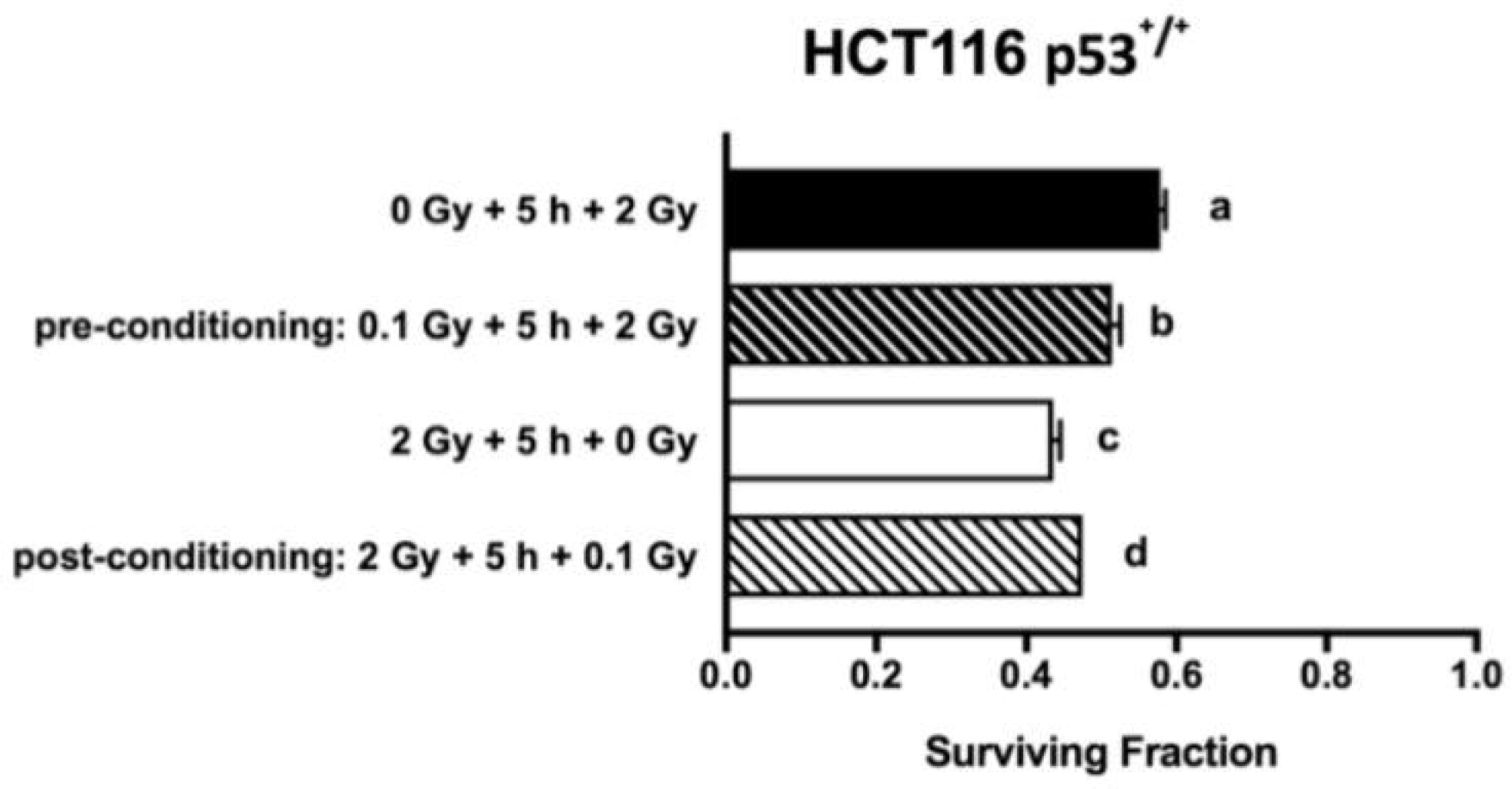
Effects of varied conditions of pre-conditioning and post-conditioning on the clonogenic survival of the human colorectal carcinoma HCT116 p53+/+ cell line. Irradiation was performed 24 h after seeding 20 cells/cm2 in triplicate T25 flasks. The priming dose was 0.1 Gy and the challenge dose was 2 Gy. The time interval between the priming and challenge dose was 5 h. The data are shown as survival fraction ± SEM (n=12) per treatment group. Based on the one-way ANOVA with Tukey’s post-test, groups with different letters are statistically significant (p<0.05) and groups sharing the same letter are not.

**Figure 7.**
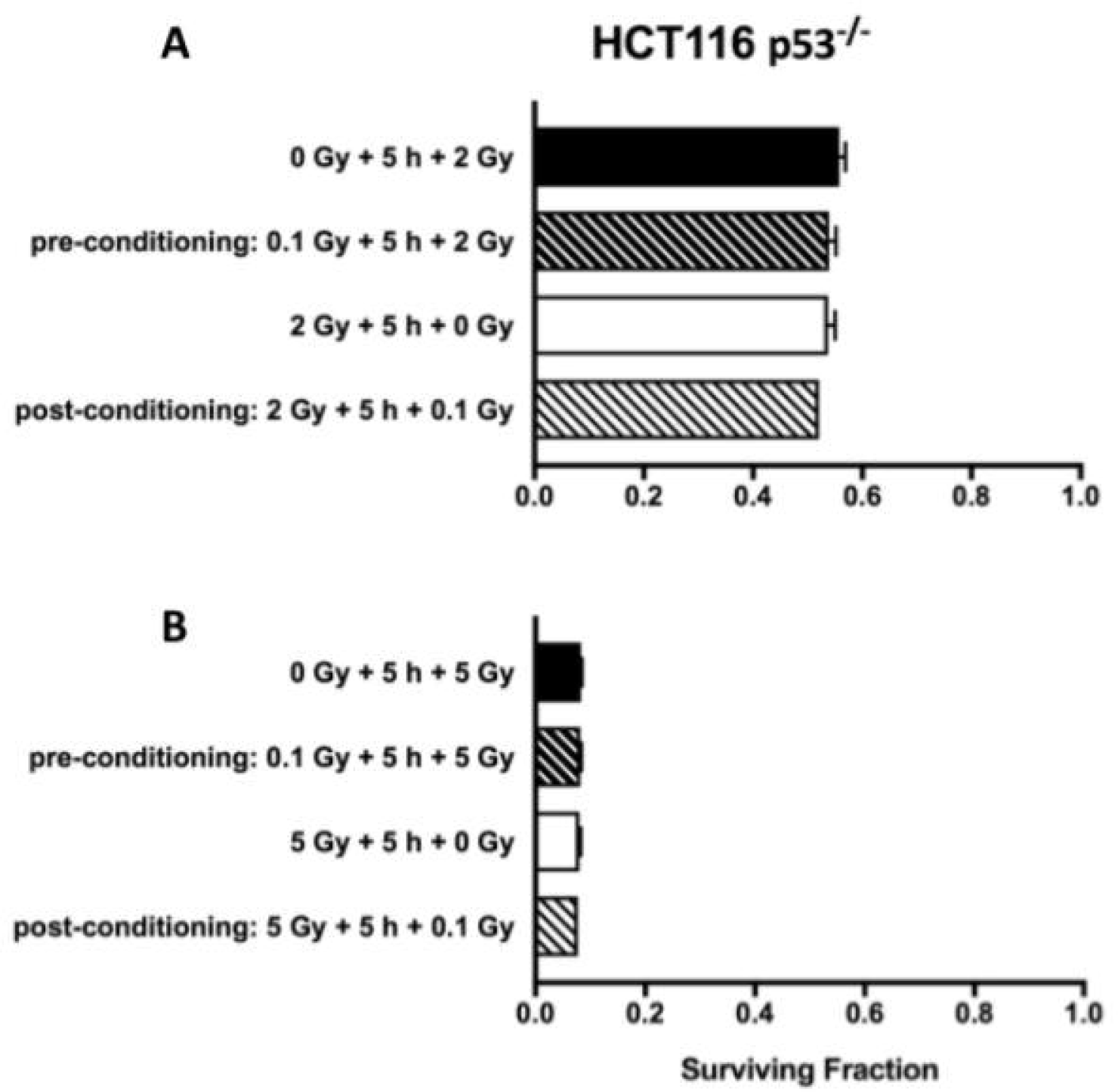
Effects of varied conditions of pre-conditioning and post-conditioning on the clonogenic survival of the human colorectal carcinoma HCT116 p53-/- cell line. Irradiation was performed 24 h after seeding 20 cells/cm2 (for 2-Gy irradiation challenge) or 60 cells/cm2 (for 4-Gy irradiation challenge) in triplicate T25 flasks. The priming dose was 0.1 Gy and the challenge dose was 2 or 5 Gy. The time interval between the priming and challenge dose was 5 h. The data are shown as survival fraction ± SEM (n=9 or n = 12) per treatment group. Based on the one-way ANOVA with Tukey’s post-test, groups with different letters are statistically significant (p<0.05) and groups sharing the same letter are not.

When the priming dose was lowered to 5 mGy and the challenge dose remained to be 2 Gy, the degree of radiosensitization by pre-conditioning was greatly reduced whereas the degree of RAR by post-conditioning was greatly enhanced in HCT116 p53^+/+^ cells (Figure 8). Interestingly, when comparing “2Gy+5h+0.1Gy” (Figure 6) to “2Gy+5h+0.005Gy” (Figure 8) in post-conditioning, the clonogenic survival was significantly higher (p<0.05, student’s t-test) when cells were primed with 5 mGy than with 0.1 Gy. This shows that 5 mGy was a more effective priming dose than 0.1 Gy in inducing post-conditioning RAR in HCT116 p53^+/+^ cells. Collectively, the data suggest that the magnitude of the priming dose was also important in dictating the cells’ radiation response outcomes.

**Figure 8.**
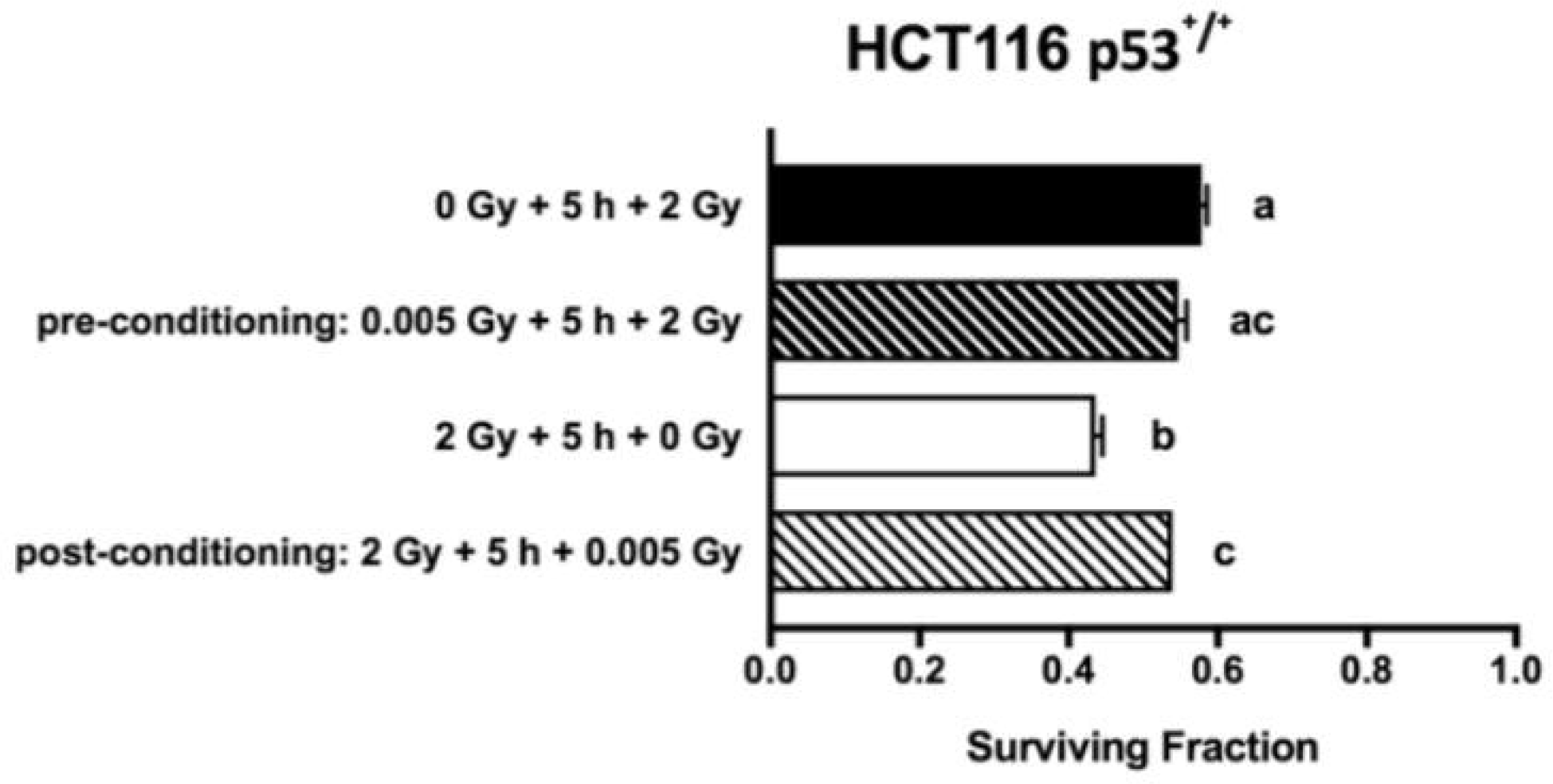
Influence of the 5 mGy priming dose on pre-conditioning and post-conditioning radiation outcomes in the human colorectal carcinoma HCT116 p53+/+ cells irradiated with a challenge dose of 2 Gy 24 h after seeding 20 cells/cm2 in triplicate T25 flasks. The time interval between the priming and challenge dose was 5 h. The data are shown as survival fraction ± SEM (n=9) per treatment group. Based on the one-way ANOVA with Tukey’s post-test, groups with different letters are statistically significant (p<0.05) and groups sharing the same letter are not.

### Correlation data between pre-/post-conditioning treatments on cell lines

For each experimental trial, cell survival from the pre-conditioning treatment was compared with the post-conditioning treatment (same priming and large challenge dose, just in reverse order) against the standard large-dose only control. The data was then plotted in Figure 9 as a difference in survival fraction between pre-/post-conditioning where the changes in survival fraction per experiment could be observed. A Pearson’s Correlation Coefficient was calculated using the 2 arrays of difference in survival fraction from pre-conditioning and difference in survival fraction from post-conditioning per experimental trial, resulting in a parallel comparison between the strength of pre-conditioning and that of post-conditioning on the cells. A positive correlation of r=0.9 was found which suggests that as there is an increase in the effects from a pre-conditioning treatment, there is similarly an increase of similar magnitude in the effects on survival fraction from post-conditioning treatment within the same trial. The data for the correlation holds the time prior to first radiation treatment constant at 6h to constrain parameters and keep experimental conditions consistent.

**Figure 9.**
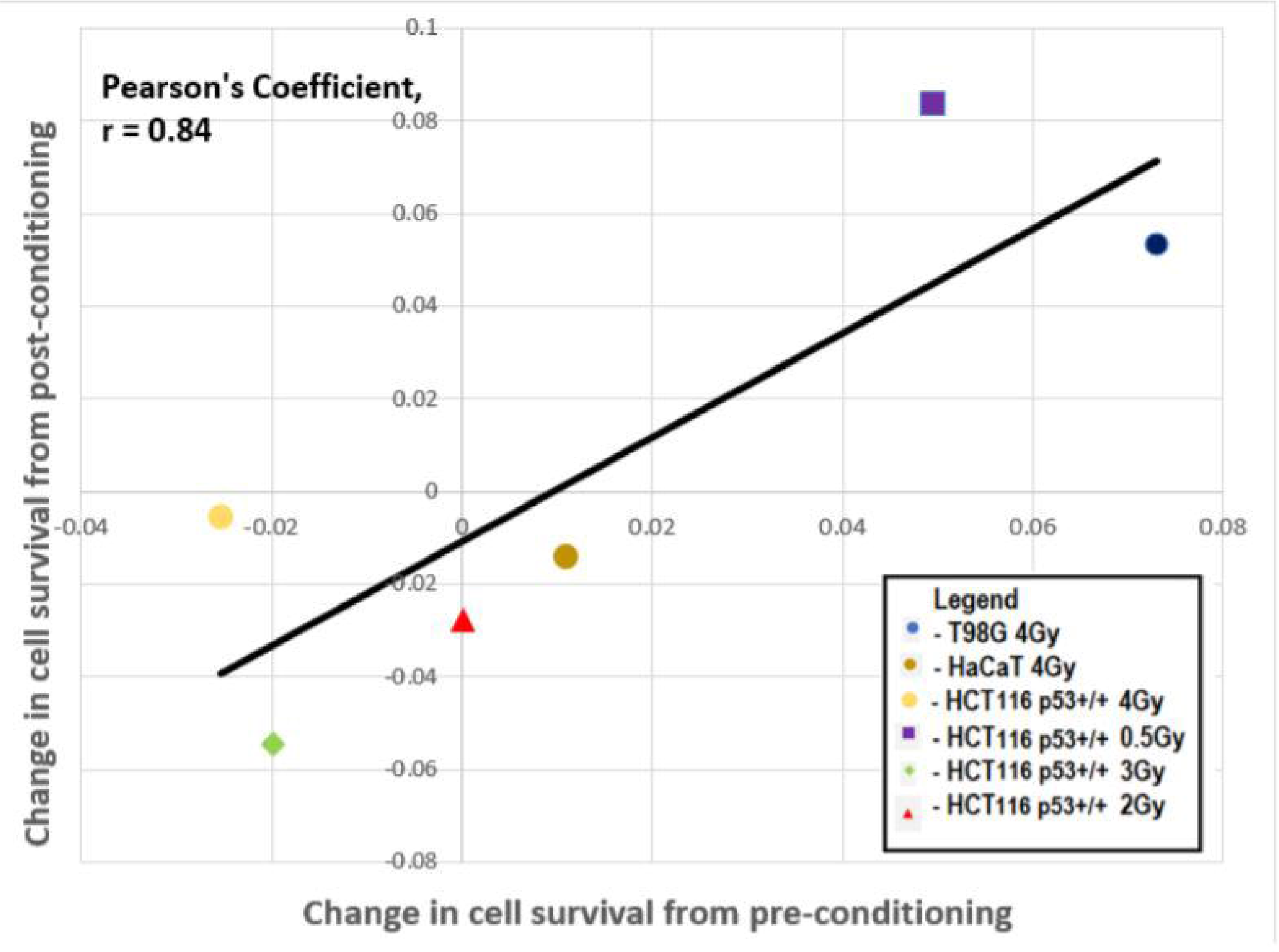
Correlation in change in cell survival after pre-conditioning and post-conditioning treatments for three cell lines (glioblastoma cells T98G, normal keratinocytes HaCaT, and colorectal carcinoma epithelial cells HCT116 p53+/+). The challenge dose was 0.5Gy, 2Gy, 3Gy, or 4Gy. Each data point represents a mean of three independent trials.

Overall, our results show that radiation conditioning responses are complex and depend on at least the following factors: the magnitude of priming/challenge dose, the time interval between priming and challenge dose, p53 status, cell seeding time prior to the first radiation treatment.

## DISCUSSION

The current study presents a unique parallel comparison between pre-conditioning and post-conditioning in different human cell types in response to gamma radiation. As radiation preconditioning had already been well established to exert an adaptive response, a particular focus in this work was to determine the consequence of post-conditioning and compare to that of pre-conditioning. The priming (smaller) and challenge (larger) doses were the same in both conditioning methods; the only difference was the opposite delivery order of the doses to delineate pre- and post-conditioning principles.

In the glioblastoma T98G cells, the observed RAR in pre-conditioning in the current work agreed with what Ryan et al. reported in 2009 using Cobalt-60 as the gamma source. The current study showed, for the first time, that the RAR could develop in glioblastoma cells when the priming dose was introduced after a large challenge dose in post-conditioning. Similar levels of increase in clonogenic survival in pre-and post-conditioning in T98G suggests that a common cellular mechanism is very likely involved in inducing protective effects induced by both conditioning methods. In general, the protective effect a result of post-conditioning was consistent with the conclusions in mouse studies by Day et al. (2007) and Lemon et al. (2017a).

In the normal keratinocyte HaCaT cells, no RAR was observed in both pre- and post-conditioning. The inability of HaCaT cells to develop an RAR in pre-conditioning also agreed with Ryan et al. (2009). The concurrent lack of RAR in both pre- and post-conditioning, in contrast to T98G, also suggests that a common cellular mechanism driving RAR may be absent in HaCaT cells. The observed differential outcomes perhaps mirror some of the results by Liu and Wu (2015). In one instance, these authors found that post-conditioning had no effect in the hamster lung fibroblast V79S cell line but had a protective effect in the human intestinal carcinoma HT-29 cell line.

In the colorectal carcinoma HCT116 p53^+/+^ cells, cellular responses after pre- and post-conditioning were complex. No RAR was observed in pre-conditioning at either the priming dose of 5 mGy or 0.1 Gy. This result was surprising because (1) cells displaying the HRS/IRR trait could develop RAR (Ryan et al., 2009) and HCT116 p53^+/+^ had HRS characteristics (Enns et al., 2004), and (2) 5 mGy had been shown to be effective in inducing pre-conditioning RAR in HCT116 p53^+/+^ recently by Murley et al. (2017). As Murley et al. (2017) used a high dose rate while the lower dose rate was used in this study, dose rate may be a differentiating factor in eliciting the RAR induction. In the present study, a protective effect was observed in post-conditioning at both priming doses of 5 mGy or 0.1 Gy. In contrast to how T98G and HaCaT cells responded to radiation conditioning, HCT11 p53^+/+^ did not develop RAR in preconditioning but did develop RAR in post-conditioning.

The noted difference between the “0Gy+5h+2Gy” and “2Gy+5h+0Gy” clonogenic survival was a surprising observation. The “0Gy+5h+2Gy” flasks were at 29 h post seeding prior to irradiation and the “2Gy+5h+0Gy” clonogenic flask were at 24 h post seeding prior to irradiation. It is possible that there were more cells that doubled at 29 h than those at 24 h. Therefore, we performed two additional experiments to see if this was a possible reason. In the first experiment, we seeded 1 million cells in T25 flasks and counted the total cell numbers in the flasks 24 or 29 h post seeding. We found that the cell numbers at 24 and 29 h were statistically indifferent from each other (Figure S1). As the cell status can be influenced under an extremely low-cell-density environment, in the second experiment we seeded cells at the clonogenic cell density in T25 flasks, stained the flasks with the fuchsin carbol dye 24 or 29 h post seeding, and microscopically scored the number of singlets (single cells) and doublets (cells that already doubled). We found that the singlet percent was the same at 24 and 29 h and the doublet percent was also the same at 24 h and 29 h (Figure S2). Therefore, observed differences in the clonogenic survival cannot be explained by the unmatched percentages of cells that have or have not divided at 24 h and 29 h. It remains possible that the observed differences could be attributable to a cell synchronization effect. However, while the 2Gy dose could arrest cells, as long as the pre- or post-conditioning flasks are assessed relative to the correct control (24 or 29 hrs), this should not be an issue. It could however provide an explanation for a post-conditioning protective effect if the cell synchronisation led to more cells being in a radioresistant cell cycle phase at the time the conditioning dose was applied. Since the conditioning dose is very small it is unlikely to cause the effect seen but the possibility is presented here for consideration. We consider it more likely that parallel processes are induced by small and large doses and that the precise order of administration of the high and low doses are less important than the absolute fact of two-dose administration. On the other hand, differences seen in cells seeded 6 h or 24 h before irradiation could be due to cell cycle arrest, synchronization or “lag” after trypsinization.

T98G and HaCaT have a mutant p53 whereas HCT116 p53^+/+^ cells have a wild-type p53 and the different p53 status could influence the outcome of RAR. Indeed, Miller et al. (2016) and Marley et al. (2017; 2018) tested a collection of mammalian cell lines with different p53 status and found that wild-type p53 cells (including HCT116 p53^+/+^) developed RAR whereas mutant or null p53 cells developed a radiosensitive response instead. In the current study, we also found that HCT116 p53^+/+^ with a wild-type p53 developed RAR with the priming dose of 5 mGy as well as 0.1 Gy. We also found that when cells (HCT116 p53^-/-^) had no p53 expression, RAR disappeared but was not replaced with a radiosensitizing response as previously reported by Miller et al. (2016) and Marley et al. (2017; 2018). On the other hand, in contrast to those authors’ work, T98G and HaCaT with a mutant p53 also did not develop radio-sensitizing response. In T98G cells, p53 has a Met237 mutation (Van Meir et al., 1994) and has no functional activities (Schmidt et al., 2001). HaCaT has His179 and Arg282 mutations (Lehman et al., 1993) and has partial functions. Arg282 is one of the hotspot mutations and frequently mutated in many cancers (Kamaraj and Bogaerts, 2015). Arg282 mutation has been shown to acquire a gain of function (Zhang et al., 2016) and is attributable to an alternative radio-responsive apoptosis pathway in HaCaT cells (Henseleit et al., 1997). The likely contribution of different p53 mutations to the different responses of cells to low-dose biophoton radiation was recently linked by Le et al. (2017b). Our results suggest that there are other factors besides p53 that regulate RAR.

The mutual relationship between HRS/IRR and RAR phenomena has been previously implicated (Marples and Joiner 1995; Ryan et al., 2009). T98G and HCT116 p53^+/+^ were HRS/IRR-positive (Short et al 1999; Enns et al., 2004) and developed RAR whereas HaCaT and HCT116 p53^-/-^ were HRS/IRR-negative (Enns et al., 2004; Ryan et al., 2009) and did not develop RAR. Although common molecular players that participate in the orchestration of mechanisms that enable HRS/IRR and RAR are not known, our data in this work further corroborated the previous notion that both phenomena may share some interconnected biochemical pathways activated by low-dose radiation.

The most surprising finding in the current study was with HCT116 p53^+/+^. Pre-conditioning had either no effect of radiosensitizing effect while post-conditioning could induce both radiosensitizing and radioresistant traits and the difference in outcomes was due to how the conditioning conditions were formulated. Our results have implications in clinical oncology where diagnostic imaging doses may present a potential challenge affecting the treatment outcome for cancers with HRS/IRR traits as well as in health physics concerning occupationally exposed nuclear workers or bystander residents living in radioactive-contaminated areas.

## ACKNOWLEDGEMENTS

The authors acknowledge support from the Canada Research Chairs program, the National Science and Engineering Research Council (NSERC) Collaborative Research and Development program and NSERC Discovery Grants program, the National CFIDS (Chronic Fatigue and Immune Deficiency Syndrome) Foundation Inc, Bruce Power and the CANDU Owner’s Group (COG). The authors kindly acknowledge Scott McMaster (McMaster University) for assistance with the use of the irradiation facility as well as dose uniformity analysis. The authors also acknowledge Dr. Cristian Fernandez-Palomo for his cultivation of T98G and passing this line to us.

## REFERENCES

Aimo A, Borrelli C, Giannoni A, Pastormerlo LE, Barison A, Mirizzi G, Emdin M, Passino C (2015). Cardioprotection by remote ischemic conditioning: Mechanisms and clinical evidences. World J Cardiol 7(10):621–632.

Audette-Stuart M, Yankovich T (2011). Bystander effects in bullfrog tadpoles. Radioprotection 46:S497–S502.

Azzam EI, Raaphorst GP, Mitchel RE (1994). Radiation-induced adaptive response for protection against micronucleus formation and neoplastic transformation in C3H 10T1/2 mouse embryo cells. Radiat Res 138(1 Suppl):S28–31.

Brattain MG, Fine WD, Khaled FM, Thompson J, Brattain DE (1981). Heterogeneity of malignant cells from a human colonic carcinoma. Cancer Res 41(5):1751–1756.

Bunz F, Dutriaux A, Lengauer C, Waldman T, Zhou S, Brown JP, Sedivy JM, Kinzler KW, Vogelstein B (1998). Requirement for p53 and p21 to sustain G2 arrest after DNA damage. Science 282:1497–1501.

Calabrese EJ, Bachmann KA, Bailer AJ, Bolger PM, Borak J, Cai L, Cedergreen N, Cherian MG, Chiueh CC, Clarkson TW, Cook RR, Diamond DM, Doolittle DJ, Dorato MA, Duke SO, Feinendegen L, Gardner DE, Hart RW, Hastings KL, Hayes AW, Hoffmann GR, Ives JA, Jaworowski Z, Johnson TE, Jonas WB, Kaminski NE, Keller JG, Klaunig JE, Knudsen TB, Kozumbo WJ, Lettieri T, Liu SZ, Maisseu A, Maynard KI, Masoro EJ, McClellan RO, Mehendale HM, Mothersill C, Newlin DB, Nigg HN, Oehme FW, Phalen RF, Philbert MA, Rattan SI, Riviere JE, Rodricks J, Sapolsky RM, Scott BR, Seymour C, Sinclair DA, Smith-Sonneborn J, Snow ET, Spear L, Stevenson DE, Thomas Y, Tubiana M, Williams GM, Mattson MP (2007). Biological stress response terminology: Integrating the concepts of adaptive response and preconditioning stress within a hormetic dose-response framework. Toxicol Appl Pharmacol 222(1):122–128.

Calabrese EJ, Mattson MP (2017). How does hormesis impact biology, toxicology, and medicine? Aging Mech Dis 3:13

Day TK, Zeng G, Hooker AM, Bhat M, Scott BR, Turner DR, Sykes PJ (2006). Extremely Low Priming Doses of X Radiation Induce an Adaptive Response for Chromosomal Inversions in pKZ1 Mouse Prostate. Radiat Res 166:757–766.

Day TK, Zeng G, Hooker AM, Bhat M, Scott BR, Turner DR, Sykes PJ (2007). Adaptive response for chromosomal inversions in pKZ1 mouse prostate induced by low doses of X radiation delivered after a high dose. Radiat Res 167:682–692.

de Toledo SM, Asaad N, Venkatachalam P, Li L, Howell RW, Spitz DR, Azzam EI (2006). Adaptive responses to low-dose/ low-dose-rate gamma rays in normal human fibroblasts: The role of growth architecture and oxidative metabolism. Radiat Res 166:849–857.

Enns L, Bogen KT, Wizniak J, Murtha AD, Weinfeld M (2004). Low-Dose Radiation Hypersensitivity Is Associated With p53-Dependent Apoptosis. Mol Cancer Res 2(10):557–566.

Fernandez-Palomo C, Seymour C, Mothersill C (2016). Inter-relationship between low-dose hyper-radiosensitivity and radiation-induced bystander effects in the human T98G glioma and the epithelial HaCaT cell line. Radiat Res 185(2):124–133.

Henseleit U, Zhang J, Wanner R, Haase I, Kolde G, Rosenbach T (1997). Role of p53 in UVB-induced apoptosis in human HaCaT keratinocytes. J Invest Dermatol 109:722–727.

Kamaraj B, Bogaerts A (2015). Structure and Function of p53-DNA Complexes with Inactivation and Rescue Mutations: A Molecular Dynamics Simulation Study. PLoS One 10(8):e0134638.

Le M, Fernandez-Palomo C, McNeill FE, Seymour CB, Rainbow AJ, Mothersill CE (2017a). Exosomes are released by bystander cells exposed to radiation-induced biophoton signals: Reconciling the mechanisms mediating the bystander effect. PLoS One 12(3):e0173685.

Le M, Mothersill CE, Seymour CB, Rainbow AJ, McNeill FE (2017b). An Observed Effect of p53 Status on the Bystander Response to Radiation-Induced Cellular Photon Emission. Radiat Res 187(2):169–185.

Lehman TA, Modali R, Boukamp P, Stanek J, Bennett WP, Welsh JA, Metcalf RA, Stampfer MR, Fusenig N, Rogan EM, Hurries CC (1993). p53 mutations in human immortalized epithelial cell lines. Carcinogenesis 14:833–839.

Lemon JA, Phan N, Boreham DR (2017a). Multiple CT scans extend lifespan by delaying cancer progression in cancer-prone mice. Radiat Res 188(4.2):492–504.

Lemon JA, Phan N, Boreham DR (2017b). Single CT Scan Prolongs Survival by Extending Cancer Latency in Trp53 Heterozygous Mice. Radiat Res 188(4.2):505–511.

Lin PS, Wu A (2005). Not all 2 Gray radiation prescriptions are equivalent: Cytotoxic effect depends on delivery sequences of partial fractionated doses. Int J Radiat Oncol Biol Phys 63(2):536–544.

Maguire P, Mothersill C, McClean B, Seymour C, Lyng FM (2007). Modulation of radiation responses by pre-exposure to irradiated cell conditioned medium. Radiat Res 167(4):485–492.

Marples B, Joiner MC (1995). The elimination of low-dose hypersensitivity in Chinese hamster V79-379A cells by pretreatment with X rays or hydrogen peroxide. Radiat Res 141(2):160–169.

Matsumoto H, Hayashi S, Hatashita M, Ohnishi K, Shioura H, Ohtsubo T, Kitai R, Ohnishi T, Kano E (2001). Induction of radioresistance by a nitric oxide-mediated bystander effect. Radiat Res 155:387–396.

Matsumoto H, Hayashi S, Hatashita M, Shioura H, Ohtsubo T, Kitai R, Ohnishi T, Yukawa O, Furusawa Y, Kano E (2000). Induction of radioresistance to accelerated carbon-ion beams in recipient cells by nitric oxide excreted from irradiated donor cells of human glioblastoma. Int J Radiat Biol 76:1649–1657.

Matsumoto H, Takahashi A, Ohnishi T (2007). Nitric oxide radicals choreograph a radioadaptive response. Cancer Res 67:8574–8579.

Miller RC, Murley JS, Rademaker AW, Woloschak GE, Li JJ, Weichselbaum RR, Grdina DJ (2016). Very low doses of ionizing radiation and redox associated modifiers affect survivin-associated changes in radiation sensitivity. Free Radic Biol Med 99:110–119.

Mitchel RE, Jackson JS, McCann RA, Boreham DR (1999). The adaptive response modifies latency for radiation-induced myeloid leukemia in CBA/H mice. Radiat Res 152:273–279.

Mothersill C, Bristow RG, Harding SM, Smith RW, Mersov A, Seymour CB (2011). A role for p53 in the response of bystander cells to receipt of medium borne signals from irradiated cells. Int J Radiat Biol 87(11):1120–1125.

Mothersill C, Seymour CB (1993). Recovery of the radiation survival curve shoulder in CHO-KI, XRS-5 and revertant XRS-5 populations. Mutat Res 285(2):259–266.

Mothersill C, Smith R, Wang J, Rusin A, Fernandez-Palomo C, Fazzari J, Seymour C (2018). Biological Entanglement-Like Effect After Communication of Fish Prior to X-Ray Exposure. Biological Entanglement-Like Effect After Communication of Fish Prior to X-Ray Exposure. Dose Response 16(1): 1559325817750067.

Murley J, Miller R, Weichselbaum R, Grdina D (2017). TP53 Mutational Status and ROS Effect the Expression of the Survivin-Associated Radio-Adaptive Response. Radiat Res 188(5):579–590.

Murley JS, Arbiser JL, Weichselbaum RR, Grdina DJ (2018). ROS modifiers and NOX4 affect the expression of the survivin-associated radio-adaptive response. Free Radic Biol Med 123:39–52.

Ryan L, Joiner M, Seymour C, Mothersill C (2009). Radiation-induced adaptive response is not seen in cell lines showing a bystander effect but is seen in lines showing HRS/IRR response. Int J Radiat Biol 85(1):87–95.

Ryan LA, Seymour CB, O’Neill-Mehlenbacher A, Mothersill CE (2008). Radiation-induced adaptive response in fish cell lines. J Environ Radioact 99(4):739–747.

Short S, Mayes C, Woodcock M, Johns H, Joiner MC (1999). Low dose hypersensitivity in the T98G human glioblastoma cell line. Int J Radiat Biol 75:847–855.

Smith D, Raaphorst G (2003). Adaptive responses in human glioma cells assessed by clonogenic survival and DNA strand break analysis. Int J Radiat Biol 79:333–339.

Smith RW, Mothersill C, Hinton T, Seymour CB (2011). Exposure to low level chronic radiation leads to adaptation to a subsequent acute X-ray dose and communication of modified acute X-ray induced bystander signals in medaka (Japanese rice fish, Oryzias latipes). Int J Radiat Biol 87(10):1011–1022.

Olivieri G, Bodycote J, Wolff S (1984). Adaptive response of human lymphocytes to low concentrations of radioactive thymidine. Science 223:594–597.

Popanda O, Zheng C, Magdeburg JR, Buttner J, Flohr T, Hagmuller E, Thielmann HW (2000). Mutation analysis of replicative genes encoding the large subunits of DNA polymerase alpha and replication factors A and C in human sporadic colorectal cancers. Int J Cancer 86:318–324.

Puck T, Marcus P (1956). Action of X-rays on mammalian cells. J Exp Med 103:653–666.

Schmidt F, Rieger J, Wischhusen J, Naumann U, Weller M (2001). Glioma cell sensitivity to topotecan: the role of p53 and topotecan-induced DNA damage. Eur J Pharmacol 2001 412(1):21–25.

Takahashi A (2001). Different inducibility of radiation- or heat-induced p53-dependent apoptosis after acute or chronic irradiation in human cultured squamous cell carcinoma cells. Int J Radiat Biol 277:215–224.

Tang H, Chen L, Liu J, Shi J, Li Q, Wang T, Wu L, Zhan F, Bian P (2016). Radioadaptive Response for Reproductive Cell Death Demonstrated in In Vivo Tissue Model of Caenorhabditis elegans. Radiat Res 185(4):402–410.

Van Meir EG, Kikuchi T, Tada M, Li H, Diserens AC, Wojcik BE, Huang HJ, Friedmann T, de Tribolet N, Cavenee WK (1994). Analysis of the p53 gene and its expression in human glioblastoma cells. Cancer Res 54(3):649–652.

Vo NTK, Sokeechand BSH, Seymour CB & Mothersill CE (2017a). Characterizing responses to gamma radiation by a highly clonogenic fish brain endothelial cell line. Environ Res 156:297–305.

Vo NTK, Sokeechand BSH, Seymour CB, Mothersill CE (2017b). Influence of chronic low-dose/dose-rate high-LET irradiation from Radium-226 in a human colorectal carcinoma cell line. Environ Res 156:697–704.

Wolff S (1998). The adaptive response in radiobiology: evolving insights and implications. Environ Health Perspectives 106(1):277–283.

Zhang Y, Coillie SV, Fang JY, Xu J (2016). Gain of function of mutant p53: R282W on the peak? Oncogenesis 5:e196.

